# Are we serologically prepared against an avian influenza pandemic and could seasonal flu vaccines help us?

**DOI:** 10.1101/2024.11.06.622244

**Authors:** Iván Sanz-Muñoz, Javier Sánchez-Martínez, Carla Rodríguez-Crespo, Corina S. Concha-Santos, Marta Hernández, Silvia Rojo-Rello, Marta Domínguez-Gil, Ahmed Mostafa, Luis Martinez-Sobrido, Jose M. Eiros, Aitor Nogales

## Abstract

The current situation with H5N1 highly pathogenic avian influenza virus (HPAI) is causing a worldwide concern due to multiple outbreaks in wild birds, poultry, and mammals. Moreover, multiple zoonotic infections in humans have been reported. Importantly, HPAI H5N1 viruses with genetic markers of adaptation to mammals have been detected. Together with HPAI H5N1, avian influenza viruses H7N9 (high and low pathogenic) stand out due to their high mortality rates in humans. This raises the question of how prepared we are serologically and whether seasonal vaccines are capable of inducing protective immunity against these influenza subtypes. An observational study was conducted in which sera from people born between years 1925-1967, 1968-1977, and 1978-1997 were collected before or after 28 days or 6 months post-vaccination with an inactivated seasonal influenza vaccine. Then, haemagglutination inhibition, viral neutralization, and immunoassays were performed to assess the basal protective immunity of the population as well as the ability of seasonal influenza vaccines to induce protective responses. Our results indicate that subtype-specific serological protection against H5N1 and H7N9 in the representative Spanish population evaluated was limited or nonexistent. However, seasonal vaccination was able to increase the antibody titers to protective levels in a moderate percentage of people, probably due to cross-reactive responses. These findings demonstrate the importance of vaccination and suggest that seasonal influenza vaccines could be used as a first line of defense against an eventual pandemic caused by avian influenza viruses, to be followed immediately by the use of more specific pandemic vaccines.

**Importance:** Influenza A viruses (IAV) can infect and replicate in multiple mammalian and avian species. Avian influenza virus (AIV) is a highly contagious viral disease that occurs primarily in poultry and wild water birds. Due to the lack of population immunity in humans and ongoing evolution of AIV, there is a continuing risk that new IAV could emerge and rapidly spread worldwide causing a pandemic, if the ability to transmit efficiently among humans was gained. The aim of this study is to analyze the basal protection and presence of antibodies against IAV H5N1, and H7N9 subtypes in the population from different ages. Moreover, we have evaluated the h humoral response after immunization with a seasonal influenza vaccine. This study is strategically important to evaluate the level of population immunity that is a major factor when assessing the impact that an emerging IAV strain would have, and the role of seasonal vaccines to mitigate the effects of a pandemic.

## 1 Introduction

Influenza A viruses (IAVs) are enveloped viruses that belong to the *Orthomyxoviridae* family. Their genomes contain eight negative-sense, single-stranded viral (v)RNA segments that are approximately 14 kb in size (1, 2). Two glycoproteins, hemagglutinin (HA) and neuraminidase (NA), are located on the viral lipid membrane, and their genetic and antigenic properties are used to classify IAVs into 18 HA (H1 to H18) and 11 NA (N1 to N11) subtypes. HA subtypes are further divided into two phylogenetic groups based on their antigenic properties: group 1 include H1, H2, H5, H6, H8, H9, H11, H12, H13, H16, H17, and H18; whereas group 2 include H3, H4, H7, H10, H14, and H15 (3–5). Viral HA and NA glycoproteins (encoded by segments 4 and 6, respectively) are also the main targets of neutralizing antibodies (NAbs) induced after vaccination and/or natural viral infection (6).

IAVs can infect and replicate in multiple mammalian and avian species and have a tremendous impact on human and animal health and the global economy. Wild aquatic birds have been considered the primary reservoir hosts for IAVs. In addition, due to IAV infections across so many different animal species, or as a consequence of viral genome reassortment, spillover infections can result in sustained epidemic transmission, when new strains with advantages in viral replication or transmission emerge in immunologically naïve populations for the newly emerging IAV strains (3–5). In addition, IAV has the potential to cause pandemics as occurred when a swine-origin H1N1 IAV (pH1N1) emerged in 2009, spreading rapidly and infecting thousands of individuals worldwide (7). In humans, IAV infections cause contagious respiratory diseases and substantial morbidity and mortality (8–10). Nowadays, two subtypes of IAVs (H1N1 and H3N2) cocirculate in humans, causing seasonal epidemics (2, 9, 11). The World Health Organization (WHO) estimates that the global disease burden from seasonal influenza results in 1 billion human infections, 3-5 million human cases of severe disease and approximately 650,000 human deaths annually (10).

Avian influenza viruses (AIVs) are highly contagious primary infect wild birds and domestic poultry. AIVs are classified in high or low pathogenic (HPAI/LPAI) depending on the molecular characteristics of the virus and its ability to cause disease and mortality in chickens. HPAIs are associated with H5- and H7-subtypes of IAV. HPAI outbreaks in poultry are no longer an occasional phenomenon in the world and the range of wild bird and mammal species affected by HPAI viruses has also expanded, with the detection of HPAI viruses frequently showing genetic markers of adaptation to mammalian hosts (3, 12–15). In addition, outbreaks of novel H5, H6, H7, H9, and H10 AIVs have caused zoonotic infections in humans and other mammals (12). Among these AIVs, the H5N1 and H7N9 subtypes stand out due to their high mortality rates in humans, which poses great public health concerns (3). Animal-to-human transmission of HPAI has occasionally occurred, with lethality rates ranging 30 −50%, while no sustainable transmission between humans has been reported (12). Important outbreaks of H5N1 HPAI infections associated sometimes with high mortality rates have been recently reported in mammals such as minks, cats, seals and dairy cattle (13, 14, 16, 17). On the other hand, LPAI and HPAI H7N9 have similar levels of morbidity and mortality in humans (4, 18, 19). Therefore, in the current scenario, H5 (HPAI) and H7 (LPAI or HPAI) AIVs pose a significant threat not only to the poultry industry but also to general public health. In fact, AIV represents an important zoonotic risk to humans, since throughout the last hundred years, avian-origin IAVs played an important role in the last four human IAV pandemics: the 1918 Spanish A/H1N1, the 1957 Asian A/H2N2, the 1968 Hong Kong A/H3N2, and the pH1N1 (4, 12, 18). In addition, AIVs have also been defined as the etiological agent of sporadic virus introductions and infections in other mammalian hosts, such dogs, and horses (5, 20, 21).

Currently, vaccination remains the main and most effective strategy to protect humans and animals against IAV infections (2, 6, 8, 11, 22). Inactivated influenza vaccines (IIVs) have been shown to be 40-60% effective in preventing morbidity and mortality associated with seasonal influenza infections (23, 24)(https://www.cdc.gov/flu/vaccines-work/effectiveness-studies.htm) by inducing humoral immune responses toward the surface viral glycoprotein HA and to less extend or in some cases to NA (25, 26). Importantly, seasonal IIVs are usually ineffective against pandemic IAV strains (18). However, previous studies have documented heterotypic responses elicited by seasonal IIVs that have led to the development of antibodies able to recognize different AIVs. Nevertheless, the extent to which seasonal IIVs can protect against heterologous subtypes in humans is unclear.

Due to the lack of population immunity in humans and the ongoing evolution of AIV, there is a continuing risk that new IAVs could emerge and rapidly spread worldwide, causing a pandemic if they gain the ability to be transmitted efficiently among humans. The aim of this study was to evaluate basal levels of protection and presence of neutralizing antibodies (NAbs) against IAV H5N1 and H7N9 subtypes in different adult age groups. Moreover, we studied the heterologous humoral response after immunization with seasonal IIV. Determining the level of pre-existing immunity in the human population, which is a major factor when assessing the impact of an emerging IAV strain, is important to evaluate the role of seasonal IIVs in mitigating the effects of a potential pandemic caused by newly introduced H5N1 and H7N9 subtypes in the human population.

## 2 Materials and Methods

### 2.1 Study design and patient recruitment

We designed a prospective study including adults and elderly people who were vaccinated with the IIV during the 2021-2022 influenza season. We recruited individuals from two different settings. The first group included workers at two automobile factories of the Renault Group in Spain in the cities of Valladolid and Palencia. The second group included individuals living in two nursing homes in the province of Valladolid (Spain), under the supervision of the “Diputación de Valladolid” Council. We selected three different age groups for this study related with the pandemics and influenza emergences of the XX century. The first group included the people born before 1968 (Group 1, G1), who were likely primed by viruses such as A/H1N1 and A/H2N2, both of which belong to group 1 IAV (27, 28). The second group included people born between 1968 and 1977 (Group 2, G2), and was likely primed by the A/H3N2 subtype, which belongs to group 2 IAV (27, 28). The third group was formed by people born after 1977 (Group 3, G3), just when the A/H1N1 re-emerged and cocirculate together with the A/H3N2, thus, any of both subtypes could have primed these people. People from G2 and G3 were only recruited from both automobile factories, and G1 was only recruited from nursing homes.

A serum sample from each individual was obtained at three different time points. The first one was just before being vaccinated with the seasonal IIV (T1). After that, a new serum sample was obtained one month after vaccination (T2), and an additional sample was collected 6-months after vaccination (T3). With this study design, we aimed to determine the immunological background of antibodies against H5N1 and H7N9 subtypes, the heterotypical response elicited by seasonal vaccines, and the duration of these antibodies over a 6-month period (**Figure 1**). During this season, the quadrivalent standard-dose inactivated vaccine FluarixTetra (GSK®) was administered to G2 and G3, and the quadrivalent high-dose inactivated vaccine Efluelda (Sanofi®) was administered to the G1. The IIV administered contained the influenza A and B strains recommended by the WHO for vaccination in the 2021–2022 season for the northern hemisphere: A/Victoria/2570/2019 pH1N1; A/Cambodia/e0826360/2020 H3N2; B/Washington/02/2019 (Victoria lineage); and B/Phuket/3073/2013 (Yamagat lineage)). This work was approved by the Ethics Committee of the Eastern Health Area of Valladolid (Cod: PI 21–2442 and PI 22-2894). All participants provided written informed consent prior to vaccination and sampling. Additionally, this research was conducted according to the Declaration of Helsinki.

**Figure 1.**
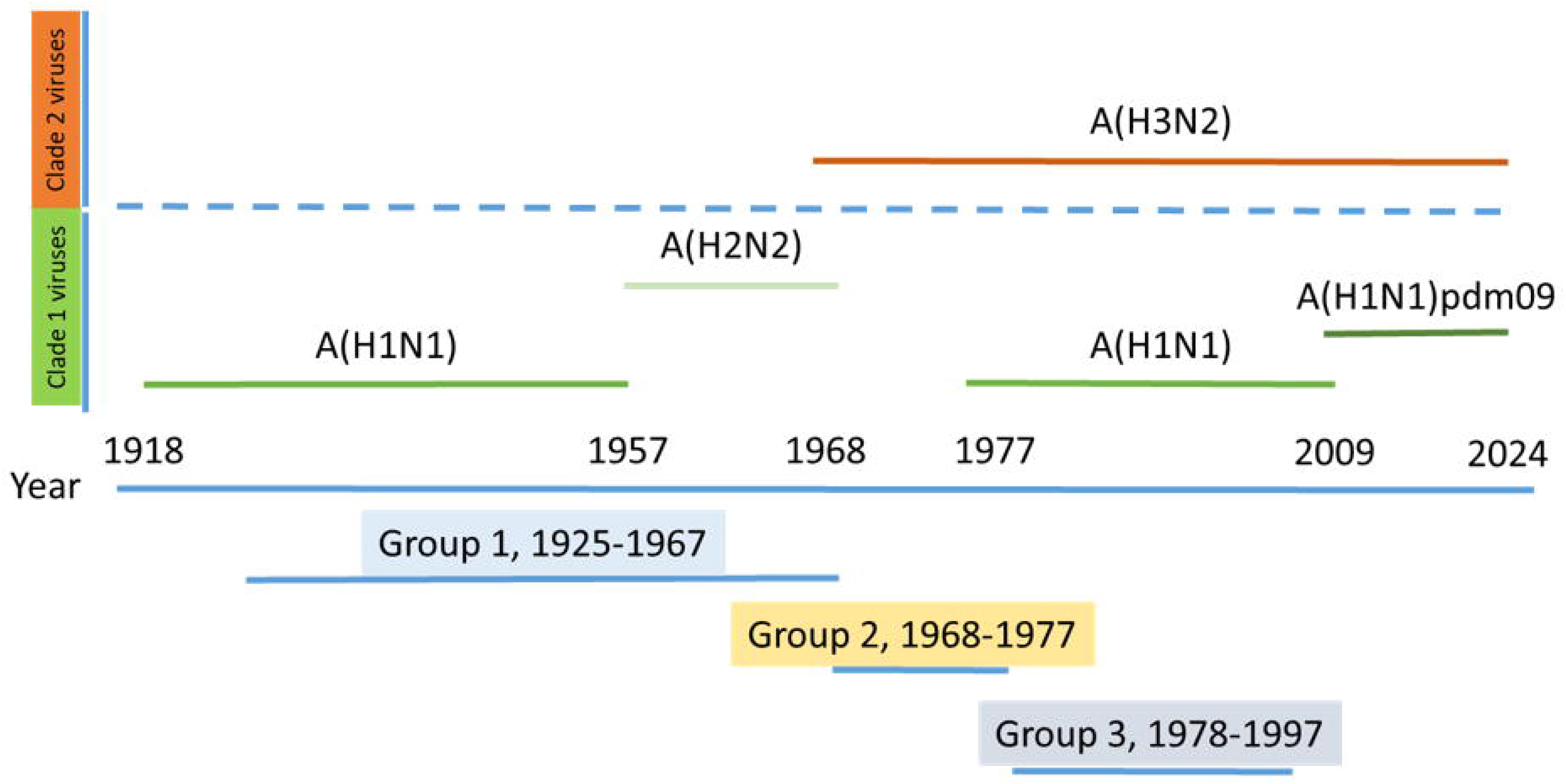
Distribution of age groups regarding the most likely imprinting by both clade 1 and 2 IAVs. The participants in the study were divided into three age groups depending on the most likely first encounter with influenza A viruses. Group 1 included people born between 1925 and 1967 who were likely primed by both the A(H1N1) and A(H2N2) clade 1 viruses. Group 2 included people born between 1968 and 1977 who were likely primed by the A(H3N2) subtype clade 2 virus. Group 3 included people born between 1978 and 1997 who were likely primed by both A(H3N2) and A(H1N1) re-emergent viruses since 1977; these groups included both clade 2 and clade 1, respectively.

### 2.2 Cell lines

Human embryonic kidney 293T (293T; ATCC CRL-11268) and Madin-Darby canine kidney (MDCK, ATCC CCL-34) cells were grown in Dulbecco’s modified Eagle’s medium (DMEM) supplemented with 5% fetal bovine serum (FBS) and 1% penicillin (100 units/ml)-streptomycin (100 μg/ml)-2 mM L-glutamine (PSG) at 37°C in air enriched with 5% CO2.

### 2.3 Culture of influenza viruses in eggs

For the serological assays, the reference viruses A/Victoria/2570/2019 pH1N1 and A/Cambodia/e0826360/2020 H3N2 of the influenza vaccine were cultured in embryonated chicken eggs to obtain viral stock to be used in the analyses. Briefly, a 1/10 pre-dilution of each virus was made in PBS and 100 µl was inoculated into the allantoic cavity of 9–11 days old embryos with a 22 G syringe (29). The inoculated eggs were incubated for 48 h at 37°C, and after that they were euthanized at 4°C for 24 h. The top of the eggshell was then removed and the allantoic fluid was collected with a pipette. This liquid was stored at 4°C for 24 hours prior to being titrated using a hemagglutination assay with chicken blood, and then labeled and stored at −80°C until use.

### 2.4 Rescue of recombinant viruses and viral infections

Recombinant IAVs were rescued using previously described ambisense reverse genetics approaches (11, 21, 30, 31). Briefly, cocultures (1:1) of 293T/MDCK cells (6-well plate format, 10^6^ cells/well, duplicates) were co-transfected, using LPF3000 (Invitrogen), in suspension with six PR8 ambisense pHW-PB2, -PB1, -PA, -NP, -M, and -NS WT or Nluc plasmids and the pHW-HA and -NA from A/chicken/Egypt/F71-F114C/2022 (H5N1, HPAIV clade 2.3.4.4b) or A/Anhui/1/2013 (H7N9, LPAIV). The H5 polybasic cleavage site motif was changed to a monobasic cleavage site. The recovered viruses were plaque purified and propagated on MDCK cells at 37°C. After viral infection, the cells were maintained in DMEM supplemented with 0.3% BSA, 1% PSG, and 1 μg/mL tosyl-sulfonyl phenylalanyl chloromethyl ketone (TPCK)-treated trypsin (Sigma) (11, 21, 30, 31). Experiments involving infectious viruses were performed at the biosafety level 3 (BSL3) facilities of the Animal Health Research Center (CISA-INIA-CSIC), Madrid (Spain).

### 2.5 Virus inactivation using beta-propiolactone

Beta-propiolactone (BPL) was combined with 10 mL of virus suspension (1 x 10^7^ PFU) and 0.5 mL of NaHCO_3_ (7.5%) to attain a final concentration of 0.05%. This buffered BPL/virus combination was then thoroughly vortexed to form a homogenous mixture and kept at 4°C for 16 h in the refrigerator. The preparation was subsequently incubated at 37°C for 2 h for hydrolysis of BPL (32, 33). After completion of treatment, the virus aliquots were stored at −80°C until further use and inactivation was confirmed via three blind-passages.

### 2.6 Virus growth kinetics

Virus replication was evaluated via a multicycle growth kinetics. Confluent MDCK cells (12-well plate format, 5×10^5^ cells/well, triplicates) were infected at an MOI of 0.001. After 1 h of virus adsorption, the cells were overlaid with DMEM containing 0.3% BSA and TPCK-treated trypsin and incubated at 37°C. At the indicated times post-infection (p.i.) (24, 48, and 72 h), the virus titers in the culture supernatants were determined by immunofocus assay (fluorescent-forming units [FFU] per milliliter) using an influenza virus monoclonal antibody (MAb) against the viral nucleoprotein, NP (HB-65; ATCC H16-L10-4R5) as previously described (15, 21, 30, 31). In addition, the presence of NLuc in the culture supernatants of cells infected with Nluc-expressing viruses was quantified using Nano-Glo luciferase substrate (Promega) following the manufacturer’s specifications and a FLUOstar Omega microplate reader (BMG Labtech).

### 2.7 Protein gel electrophoresis and Western-blot analysis

Cell extracts from either mock- or virus-infected (MOI, 0.01) MDCK cells (6-well plate format, 10^6^ cells/well) were lysed at 24 h p.i. in radioimmunoprecipitation assay (RIPA) buffer, and proteins were separated by denaturing electrophoresis. The membranes were blocked for 1 h with 5% dried skim milk in PBS containing 0.1% Tween 20 (T-PBS) and incubated overnight at 4°C with specific primary monoclonal or polyclonal antibodies (pAbs): NP (MAb HB-65; ATCC H16-L10-4R5), NLuc (MAb, R&D Systems), HA (goat pAb NR-2705 for H5, goat pAb NR-48597 for H7, BeiResources), and actin (MAb A1978; Sigma), which was used as an internal loading control. Bound primary antibodies were detected with horseradish peroxidase (HRP)-conjugated secondary antibodies against the different species (Sigma). Proteins were detected by chemiluminescence (Thermo Fisher Scientific) following the manufacturer’s recommendations and photographed using a Bio-Rad ImageStation.

### 2.8 Indirect immunofluorescence assays

MDCK cells (12-well plate format, 5×10^5^ cells/well) were mock infected or infected (MOI, 0.01) with the indicated viruses. At 24 h p.i., the cells were fixed with 4% paraformaldehyde (PFA) and permeabilized with 0.5% Triton X-100 in PBS for 15 min at room temperature. Immunofluorescence assays were performed using primary HA (goat pAb NR-2705 for H5, goat pAb NR-48597 for H7, BeiResources), NA (goat pAb NR-9598 for H5, rabbit pAb NR-49276 for H7, BeiResources), and NP (MAb HB-65) antibodies and secondary Alexa-fluor (594)-conjugated antibodies against the different species. Images were taken with a Zeiss Axio fluorescence microscope (Zeiss).

### 2.9 Plaque assays and immunostaining

Confluent monolayers of MDCK cells (6-well plate format, 10^6^ cells/well) were infected with the indicated viruses for 1 h at room temperature, overlaid with agar with or without TPCK-treated trypsin, and incubated at 37°C. At 3 days p.i., the cells were fixed with 10% PFA and the overlays were removed. Then, the cells were permeabilized (0.5% Triton X-100 in PBS) for 15 min at room temperature and prepared for immunostaining as previously described (15, 21, 30, 31) using an anti-NP MAb (HB-65; ATCC H16-L10-4R5). Immunostaining was developed using vector kits (Vectastain ABC kit for mouse antibodies and DAB HRP substrate kit; Vector) following the manufacturers’ specifications.

### 2.10 Enzyme-Linked Immunosorbent Assay (ELISA)

H5- and H7-specific serum IgG antibodies were detected by ELISA. Briefly, 96-well plates were coated for 16 h at 4°C with 100 ng per well of H5 or H7 (NR-59424 and NR-44365, respectively, BeiResources) recombinant protein. After washing with PBS, the coated wells were blocked with PBS-T (Tween, 0.05%) containing 5% BSA and then incubated with a 1:50 dilution of human serum at 37°C. After 1 h of incubation, the plates were washed with PBS-T and incubated with HRP- conjugated goat anti-human IgG for 1 h at 37°C. The reactions were developed with tetramethylbenzidine (TMB) substrate (BioLegend) for 10 min at room temperature, quenched with 2 N H_2_SO_4_, and read at 450 nm (FLUOstar Omega microplate reader (BMG Labtech)).

### 2.11 Haemagglutination inhibition (HAI) assay

HAI assay was used as the reference assay to analyze the antibody response induced by influenza vaccines (34). To that end, viruses were first inactivated with BPL. Additionally, IAV strains included in the seasonal administered vaccine (H1 and H3) were used to compare the results of AIV with those of seasonal strains. Serum samples were centrifuged at 2,500 rpm for 10 min, and the supernatants were stored at −20°C. Antibody analysis against HA was performed using HAI assay. For each serum sample, 100 μl was added to 300 μl of receptor destroying enzyme (Denka Seiken, Tokyo, Japan) to remove nonspecific hemagglutination inhibitors. Following the manufacturer’s instructions, this mixture was incubated at 37°C in a water bath for 12–18 hours, followed by enzyme inactivation for 1 h at 56°C. To perform HAI, 50 µl serial 2-fold dilutions of each serum (starting dilution of 1:10) were made in 96 microwell V plates, and then 50 µl of a standard containing 4 units of hemagglutinin (HAU) was added to each well and incubated for 30 min at room temperature. Next, 50 µl of 0.5% chicken erythrocytes were added and incubated at room temperature for another 30 min. The titer of antibodies was defined as the highest dilution that completely inhibited hemagglutination. The limit of detection of the assay was 1:10. When the antisera titer was below a detectable threshold, due to a shortage or lack of antibodies, it was conventionally expressed as 5, which is half of the lowest detection threshold. The pre-vaccine, post-vaccine, and 6-month HAI titers were included in the database for study.

### 2.12 Reporter-based microneutralization (MN) assay

MN assays were performed as previously described (30, 31, 35). Briefly, human sera samples or control sera specific for anti-H5 and anti-H7 (NR-665 and NR-48597, respectively) were serially 4- fold diluted (starting dilution 1/10 for human samples) or 2-fold diluted (starting dilution 1/40 for control sera samples). Then, 200 PFU of Nluc-expressing H5N1 or H7N9 viruses were added to the serum dilutions and the mixture was incubated for 1 h at room temperature. MDCK cells (96-well plate format, 5 × 10^4^ cells/well) were infected with the serum-virus mixture and incubated for 48 h at 37°C. NLuc activity in the culture supernatants was quantified using Nano-Glo luciferase substrate (Promega) and a Lumicount luminometer (FLUOstar Omega microplate reader (BMG Labtech)). The luciferase values of virus-infected cells in the absence of antibody were used to calculate 100% viral infection. Cells in the absence of viral infection were used to calculate the luminescence background. After the Nluc concentration was measured, the cells were fixed and stained with crystal violet solution.

### 2.13 Statistical analysis

The statistical analysis was focused on a descriptive and inferential study of the antibody titers based on the age of the patients. The results were analyzed using the classical serological parameters of the European Medicines Agency (EMA) for the evaluation of vaccine efficacy (36, 37). These parameters included the seroprotection rate (SPR), seroconversion rate (SCR) and geometric mean titer (GMT). Seroprotection was considered when a HAI titer of 1/40 was achieved (38), and seroconversion was defined as a titer increase of at least fourfold between pre- and post-vaccination sera. In addition, seroconversion was considered to have occurred in cases of negative pre-vaccination sera that achieved 1/40 titers after vaccination. The values are presented as the means (95% CIs). The volunteer characteristics, SPR and SCR were compared using χ2 tests. GMTs were compared using Student’s t-test. The data from the neutralization assays were compared using two-way ANOVA. Data analysis was performed using GraphPad Prism and Microsoft Excel. A P value <0.05 was considered to indicate statistical significance.

## 3 Results

### 3.1 Patients features

A total of 135 individuals were recruited for this study: 41 were included in G1 (31.5%), 49 in G2 (37.7%), and 40 in G3 (30.8%). The mean age was 79.4 (95% CI, 74.8-84.1) years for G1, 48.4 (95% CI, 47.5-49.2) years for the G2, and 35.1 (95% CI, 33.4-36.8) years for G3. There were 22 males in G1 (53.7%), 24 males in G2 (48.9%), and 18 males in G3 (45.0%) (**Table 1**).

**Table 1.** Cohort description and epidemiological characteristics.

|  | Group 1 (1925-1967) | Group 2 (1968-1977) | Group 3 (1978-1997) |
| --- | --- | --- | --- |
| **N° (%)** | 41 (31.5) | 49 (31.5) | 40 (30.8) |
| **Age, Mean (CI95%)** | 79.4 (74.8-84.1) | 48.4 (47.5-49.2) | 35.1 (33.4-36.8) |
| **Men, n (%)** | 22 (53.7) | 24 (48.9) | 18 (45.0) |
| **Type of Vaccine** | Quadrivalent High-Dose Inactivated | Quadrivalent standard dose inactivated | Quadrivalent standard dose inactivated |

### 3.2 Generation and *in vitro* characterization of recombinant IAV

We used plasmid-based reverse genetics methods to generate the indicated recombinant IAV in the backbone of PR8 containing H5N1 or H7N9 viral HA and NA segments and the PR8 NS-WT or NS-Nluc segment (**Table 2**). Importantly, we removed the polybasic amino acids that are linked with high virulence from the H5 cleavage site. This was not necessary for the H7 viral segment, since a LPAI H7 was used in this study. To engineer the replication-competent reporter-expressing IAV, the sequence of Nluc was cloned in the NS segment that encodes NS1 and NEP, as we have described previously (30, 31, 39). We chose NLuc due to its small size (approximately 20 kDa), stability, and brightness (42, 50). Briefly, Nluc was cloned in frame with the NS1 sequence and silent mutations in the splice acceptor site of the NS segment were introduced to eliminate splicing (30, 31, 39). Then, the Porcine teschovirus-1 (PTV-1) 2A cleavage site was cloned between NS1 and NEP and the NS1 and NEP N-terminal overlapping region was duplicated after the PTV-1 2A site to produce a full-length NEP.

**Table 2.** Viruses generated in this study.

| Virus | Backbone | HA and NA genes | Reporter gene |
| --- | --- | --- | --- |
| PR8(H5N1) | A/Puerto Rico/8/1934 (H1N1) | A/chicken/Egypt/F71-F114C/2022 (H5N1, 2.3.4.4b), HA_Monobasic | No |
| PR8(H5N1)-Nluc |  |  | Nluc |
| PR8(H7N9) |  | A/Anhui/1/2013 (H7N9, LP) | No |
| PR8(H7N9)-Nluc |  |  | Nluc |

The identity of the recombinant IAVs generated was first confirmed by indirect immunofluorescence microscopy using specific serum against H5 (H5N1), N1 (H5N1), H7 (H7N9) or N9 (H7N9). In addition, a MAb against IAV NP was used as a control (**Figure 2A**). For that, monolayers of MDCK cells were mock infected or infected with the different viruses. Then, at 24 h p.i., the cells were fixed and viral infections were assessed. As expected, H5 and N1 were detected in cells infected with PR8(H5N1) or PR8(H5N1)-Nluc, while H7 and N9 antigens were detected only in MDCK infected with PR8(H7N9) or PR8(H7N9)-Nluc. Viral NP located in the nucleus of cells infected with all the viruses. The plaque phenotype of the recombinant viruses was also examined by plaque assays and immunostaining (**Figures 2B and C**). To confirm the monobasic cleavage site, in the case of PR8(H5N1) and PR8(H5N1)-Nluc, plaque assays were performed in the presence and absence of TPCK-trypsin (**Figure 2B**). The plaque sizes of PR8(H5N1) and PR8(H5N1)-Nluc were similar, and they were observed only in the presence of TPCK-trypsin (**Figure 2B**). However, the plaques of the PR8(H7N9) and PR8(H7N9)-Nluc infected cells were significantly smaller than those of the H5N1- infected cells (**Figure 2C**). Next, the expression of Nluc was confirmed by Western blot (**Figures 2D and E**). Total cell extracts from either mock-, or IAV-infected MDCK cells collected at 24 h p.i. were tested using antibodies specific for H5 or H7, NP, and NLuc. We used an antibody against actin as a loading control. Western blot analysis revealed specific bands with the expected molecular sizes for H0, H1 and H2 of H5 in cells infected with PR8(H5N1) and PR8(H5N1)-Nluc (**Figure 2D**) or H7 in cells infected with PR8(H7N9) and PR8(H7N9)-Nluc (**Figure 2E**). Moreover, specific bands for Nluc were detected only in cell extracts from MDCK cells infected with PR8(H5N1)-Nluc or PR8(H7N9)-Nluc (**Figures 2D and E**). As expected, NP levels were detected in all infected cells but not in mock-infected cells (**Figures 2D and E**).

**Figure 2.**
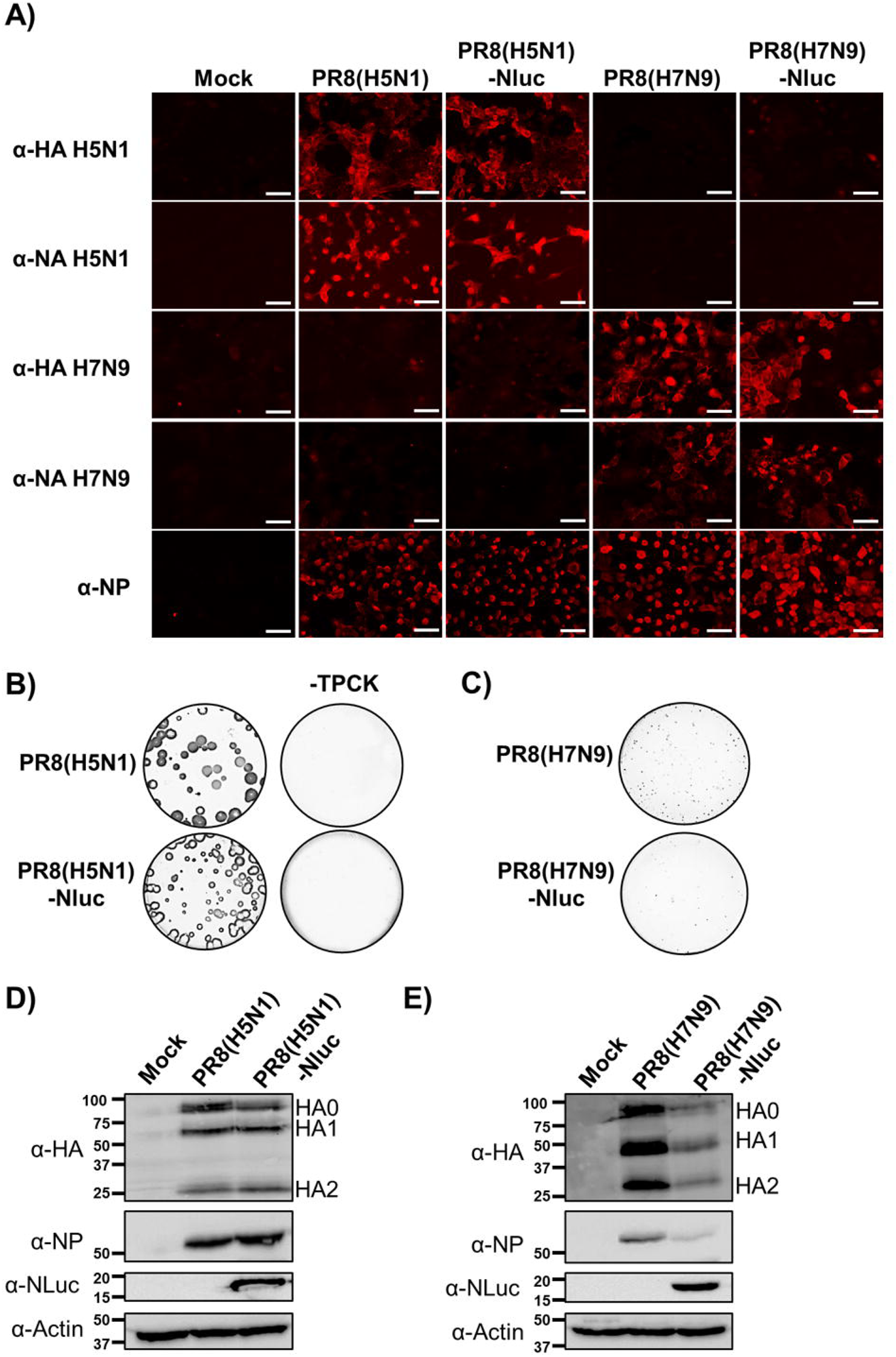
In vitro characterization of the recombinant viruses generated in this study. A) Analysis of protein expression by immunofluorescence. MDCK cells (24-well plates, 2 x 10^5^ cells/well) were infected with the indicated viruses (MOI, 0.01) or mock infected. Infected cells were fixed and permeabilized at 24 h p.i. The cells were stained using specific pAbs against H5 (NR- 2705), N1 (NR-9598), H7 (NR-48597) or N9 (NR-49276). A MAb against viral NP was used as control. Representative images are shown. Bars, 100 μm. **B and C) Plaque phenotype.** MDCK cells (6-well plate format, 1 x 10^6^ cells/well) were infected with ∼50 FFU of PR8(H5N1) or PR8(H5N1)- Nluc in the presence of absence of TPCK-trypsin (**B**) or infected with PR8(H7N9) or PR8(H7N9)- Nluc **(C)** and incubated at 37 °C for 3 days. Plaques were evaluated by immunostaining using a MAb against IAV NP (MAb HB-65). **D and E) Analysis of protein expression by western-blot.** MDCK cells (6 well plates, 10^6^ cells/well) were infected with the indicated viruses or mock infected. Protein expression was examined by Western blotting using specific antibodies against HA (NR-2705 for H5 and NR-48597 for H7), NP, and NLuc. Actin was used as a loading control. The numbers on the left indicate the molecular size of the protein markers (in kilodaltons).

The fitness of the recombinant viruses was evaluated in cell culture (**Figure 3**). To this end, confluent monolayers of MDCK cells were infected (MOI of 0.001) with PR8(H5N1) and PR8(H5N1)-Nluc or PR8(H7N9) and PR8(H7N9)-Nluc, and the presence of the virus in the culture supernatants was quantified at different h p.i. (**Figures 3A and C**). The replication of Nluc-expressing viruses was slightly delayed, and in the case of PR8(H7N9)-Nluc did not reach titers similar to those of PR8(H7N9). In addition, NLuc activity in culture supernatants was quantified at the same time points (**Figures 3B and D**). NLuc activity was detected at 24 h p.i. (the first time point assessed) and increased in a time-dependent manner, peaking at 72 h p.i. (the last time point assessed), most likely because of the Nluc release due to the cytopathic effect (CPE) caused by viral infection, as previously observed (30).

**Figure 3.**
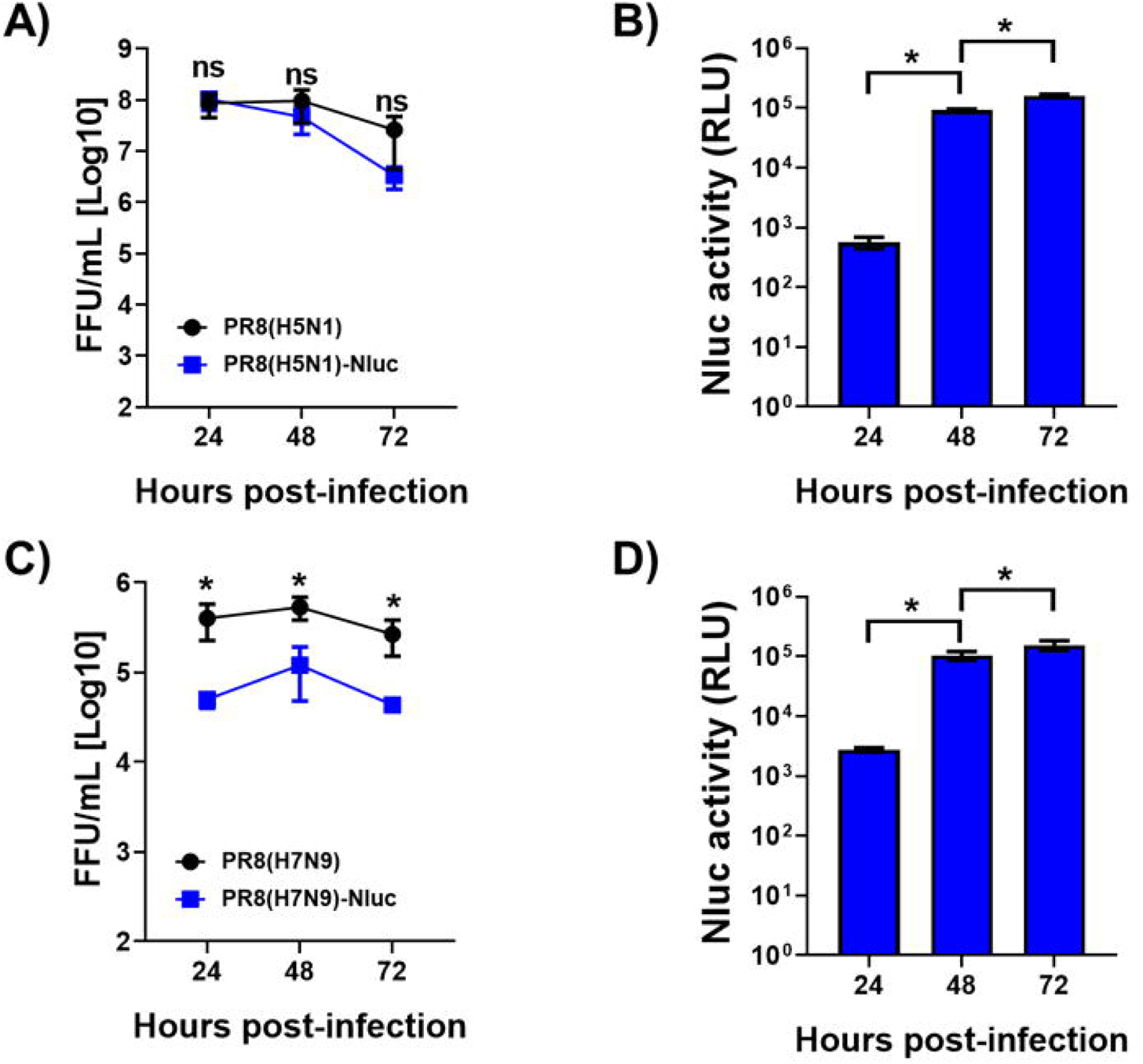
Growth kinetics of the generated recombinant viruses. (A and B) Multicycle growth kinetics. Viral titers (in FFU per milliliter) in culture supernatants from MDCK cells (12-well plates, 5 x 10^5^ cells/well, triplicates) infected with PR8(H5N1) or PR8(H5N1)-Nluc (A) or infected with PR8(H7N9) or PR8(H7N9)-Nluc **(C)** (MOI, 0.001) were determined by immunofocus assay at the indicated times post-infection. The data represent the means ± SD of triplicate samples. *, *P* < 0.05, using two-way ANOVA. **(B and D) NLuc expression.** NLuc was evaluated in the same culture supernatants obtained from the experiment, and the results are presented in panel A. RLU, relative light units.

### 3.3 Determining human immunity against IAV subtypes H5N1 and H7N9 and evolution of the response after IIV

Antibodies were first evaluated by HAI using inactivated AIV and also live IAV pH1N1 and H3N2 strains propagated in 10-day-old chicken embryonated eggs in the laboratory (**Figure 4**). Before vaccination, only 2.4% of people in G1 had protective antibodies against H5N1, and none protective antibodies were observed against H7N9 in any vaccinated group **(Figure 4A)**. Seasonal vaccination elicited heterotypic seroconversion in 15.0% of the people from the G3 and 12.2% of the people in G2, but we did not observe any response in G1 **(Figure 4B)**. The unique response observed against the H7N9 subtype was present in G2, with 2.0% of people seroconverted. In contrast, seroconversion and seroprotection rates were up to 39.0% for pH1N1 and 24.4% for A/H3N2, depending on the group. After six months (T3), the percentage of seroprotection against AIV was less than 2% in any case **(Figure 4A)**.

**Figure 4.**
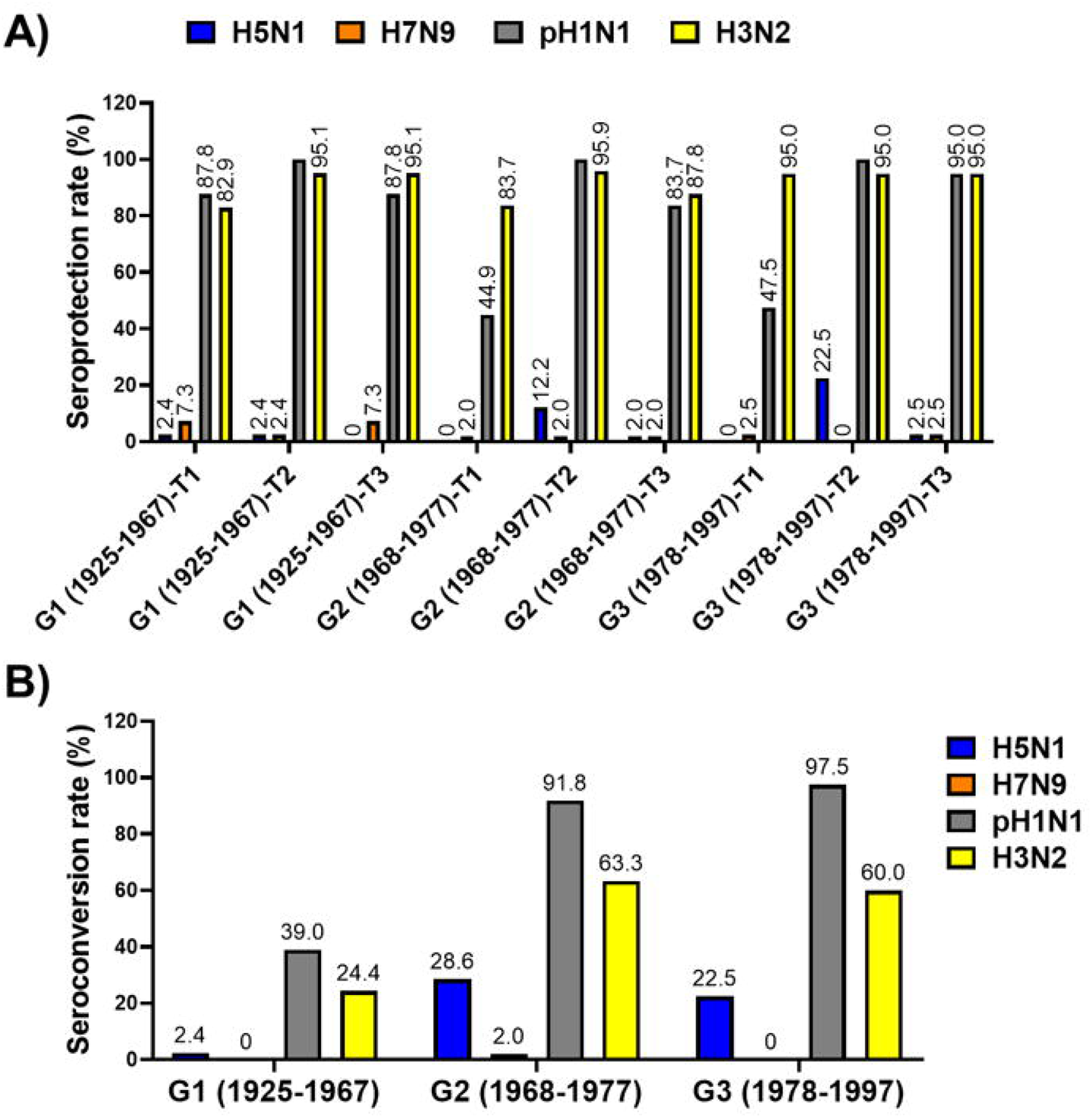
Seroprotection and seroconversion rates for IAV subtypes H1, H3, H5 and H7 in human serum samples. The seroprotection rate (**A**) and seroconversion rate (**B**) for the indicated IAV subtypes were determined in human serum samples from individuals born to 1925-1967 (G1), 1968-1977 (G2), and 1978-1997 (G3) and collected immediately before vaccine administration (T1), and post-vaccination at 28 days after vaccination (T2) or six months after vaccination (T3).

This heterotypic response after vaccination was also corroborated by the study of the geometric mean titers (GMTs) **(Figure 5)**. After vaccination with the seasonal IIV, the GMTs against the H5N1 subtype were significantly greater in G2 and G3 than in the pre-vaccine sample (p<0.01 and p<0.001 respectively). However, in the case of G1, we observed a continuous decline in the GMTs after vaccination **(Figure 5A)**. In the case of H7N9, no significant differences were detected between G2 and G3, but a significantly greater GMT was detected in G1 during T1 than during T2 (p<0.01) **(Figure 5B)**. A high to moderate decrease in the GMTs was observed for all AIV and seasonal viruses after 6 months (T3), reaching similar titers to those before vaccination for both AIVs but greater for both the H1 and H3 seasonal viruses.

**Figure 5.**
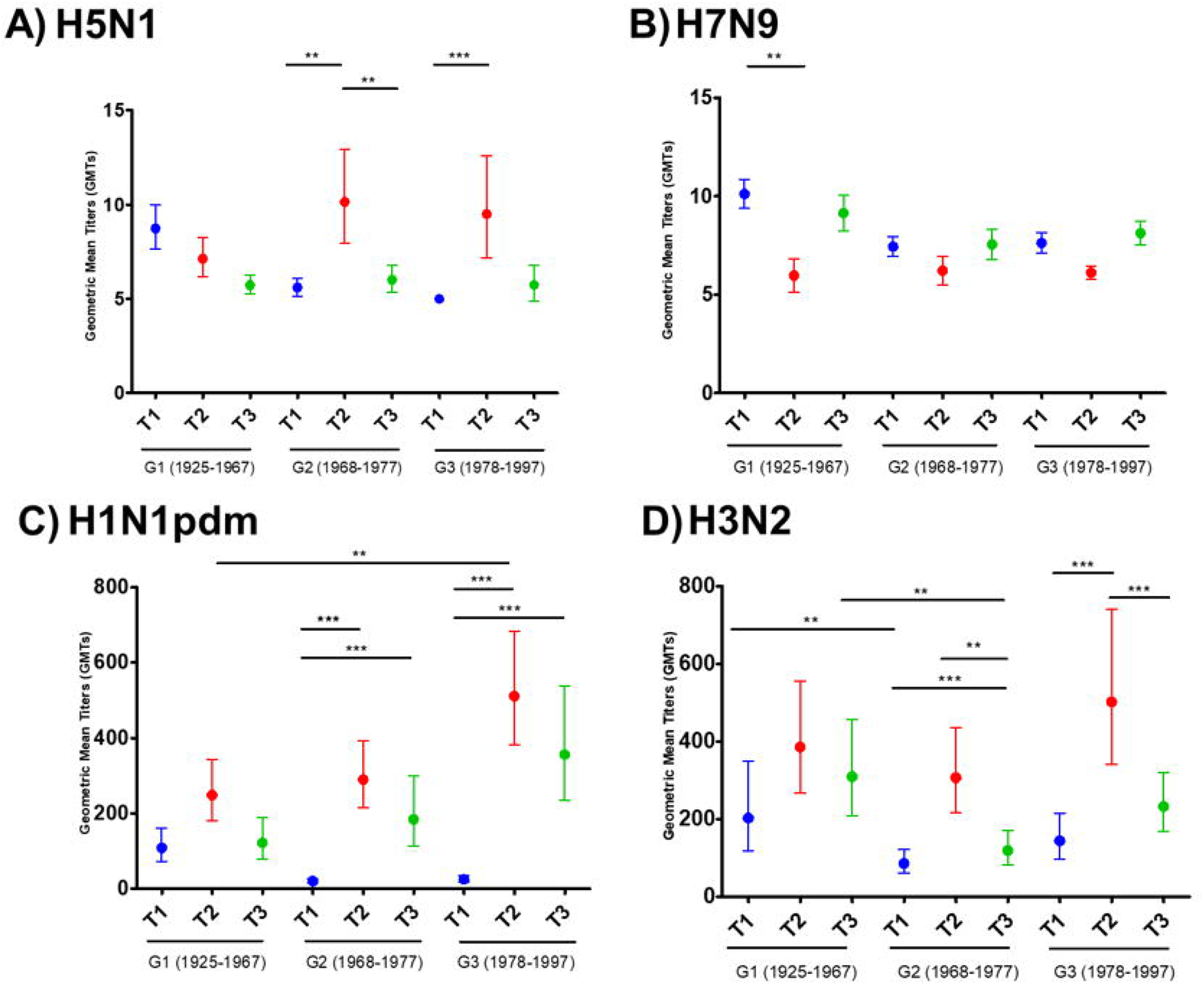
Geometric mean titers (GMTs) of hemagglutination inhibition (HAI) antibodies against IAV subtypes H1, H3, H5 and H7 in human serum samples. HAI titers against the indicated IAV subtypes were determined in human serum samples corresponding to individuals born at 1925-1967 (G1), 1968-1977 (G2), and 1978-1997 (G3) and collected immediately before vaccine administration (T1), and post-vaccination at 28 days after vaccination (T2) or six months after vaccination (T3). Statistical analysis was performed using one-way ANOVA with Bonferroni’s multiple comparison test. *\*p* < 0.05; *\*\*p* > 0.01; *\*\*\*p* < 0.001; *\*\*\*\*p* < 0.0001.

In addition, the levels of IgG antibodies recognizing H5 or H7 viral antigens were measured by ELISA using purified recombinant H5 or H7 proteins (**Figure 6**). The results indicated the presence of antibodies recognizing H5 (**Figure 6A**) and H7 (**Figure 6B**) before and after vaccination. For H5, the levels of these antibodies were significantly higher in G1 for T1 and T3, but not for T2. In addition, an increase in anti-H5 antibodies was observed in T2 for both G2 and G3. Interestingly, although high levels of anti-H7 antibodies were detected by ELISA, we did not observe significant differences for G1 or G2, and only some differences were observed for G3, where after vaccination (T2), an increase in the levels of anti-H7 antibodies was observed. Importantly, anti-H5 or anti-H7 antibodies could be recognizing conserved regions of the viral HA protein. Based on the above HAI studies, the data suggest that most likely most of the antibodies are not NAbs. In fact, since most of the population has been vaccinated against IAV, infected with IAV, or both (even multiple times), the results obtained by ELISA were, to some extent, expected.

**Figure 6.**
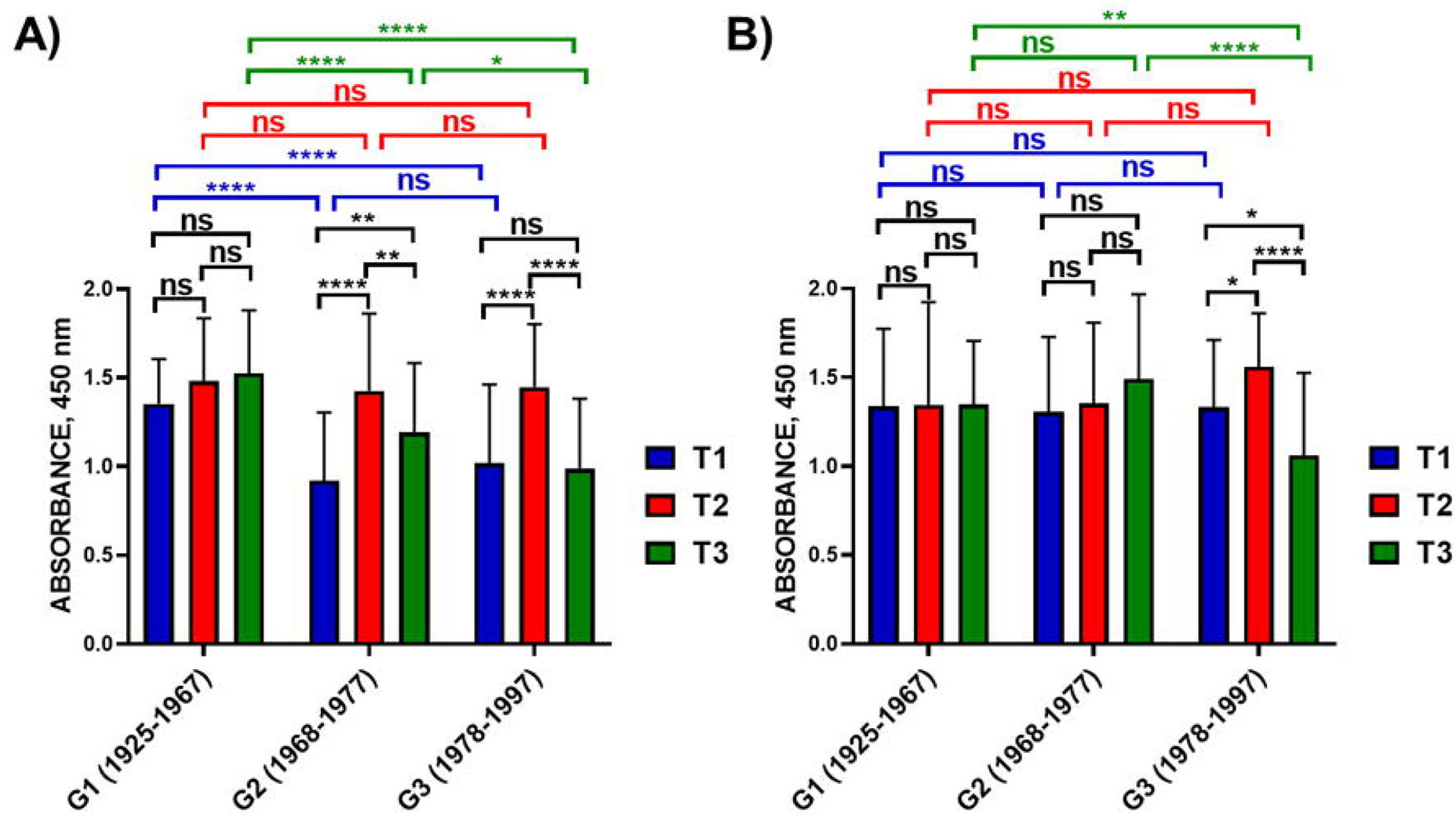
Enzyme-linked immunosorbent assay (ELISA). The presence of IgG antibodies recognizing H5 (**A**) or H7 (**B**) was evaluated by ELISA using purified recombinant H5 (NR-59424, A/bald Eagle/Florida/125/2017) or H7 (NR-44365, A/Anhui/1/2013) proteins. Human serum samples corresponding to individuals born between 1925-1967 (G1), 1968-1977 (G2), and 1978-1997 (G3) were collected immediately before vaccine administration (T1), and post-vaccination at 28 days after vaccination (T2) or six months after vaccination (T3). *, *P* < 0.05, using two-way ANOVA.

### 3.4 Cross-reactive antibodies to neutralize IAV H5N1 in vitro

NAbs are the desired immunological outcome for the induction of protective immunity after influenza vaccination or infection (24, 35, 40–42). Assess to evaluate the presence of NAbs are time-consuming and usually involve secondary methods to detect the presence of the virus. We have previously demonstrated that this limitation can be circumvented by using IAV harboring fluorescent or luciferase reporter genes whose expression can be monitored or quantified directly (30, 31, 35). To that end, we developed Nluc-expressing viruses to test the presence of NAbs in an Nluc-based MN assay. To demonstrate that PR8(H5N1)-Nluc or PR8(H7N9)-Nluc can be used to easily identify NAbs, confluent monolayers of MDCK cells were infected with each virus that had previously been incubated with H5- or H7- specific serum (NR-665 and NR-48597, respectively) (**Figure 7**). Then, at 48 h p.i. NLuc activity in the culture supernatants was quantified. As expected, PR8(H5N1)-Nluc and PR8(H7N9)-Nluc were specifically neutralized by H5- or H7-specific antibodies, respectively (**Figure 7**), demonstrating the feasibility of using PR8(H5N1)-Nluc or PR8(H7N9)-Nluc and the Nluc-based MN assay to detect NAbs.

**Figure 7.**
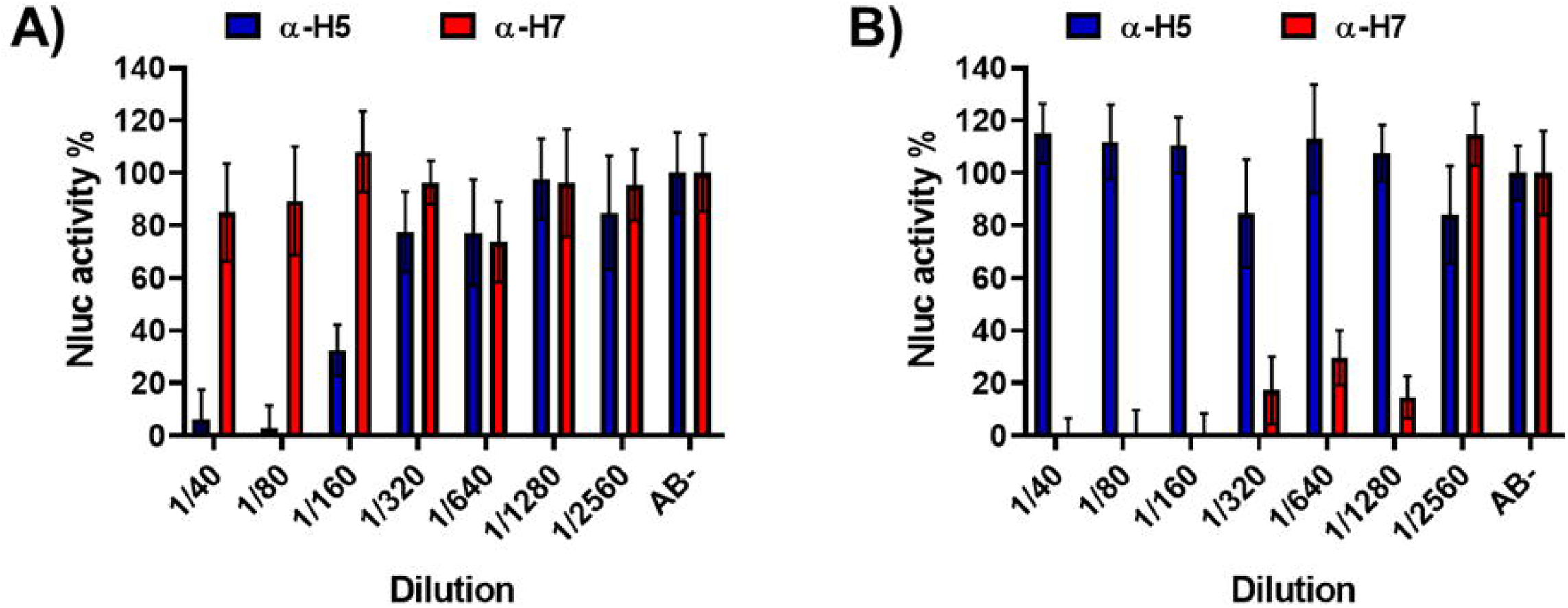
Nluc-based microneutralization assay for evaluating IAV NAbs. One-two hundred FFU of PR8(H5N1)-Nluc (**A**) or PR8(H7N9)-Nluc (**B**) viruses were pre-incubated with 2-fold serial dilutions (starting dilution 1/40) of the indicated sera for PR8 H5N1 (NR-665) or PR8 H7N9 NR- 48597) for 1 h. Subsequently, MDCK cells (96-well plates, 2 x 10^4^ cells/well, and triplicates) were infected with the antibody-virus mixture. Virus neutralization was determined by quantitating NLuc reporter expression at 48 h p.i.. Mock-infected cells were used as internal controls for basal levels of luciferase (NLuc) expression. Infected cells in the absence of serum were used to determine the maximum reporter expression. The data are presented as the means of triplicate analyses.

Next, pre-existing immunity (T1) against IAV H5N1 in 135 individuals from different groups of age (G1, G2 and G3) were tested. In addition, whether seasonal IIV can induce heterosubtipic responses and NAbs against IAV H5N1 was also determined (T2 and T3). For that, the ability of serum samples collected before (T1) or after (T2 and T3) (**Table 1**) vaccination to neutralize PR8(H5N1)- Nluc was evaluated using our Nluc-based MN assay. F**igure 8A** shows a heatmap of the neutralization levels observed for each individual sample, and the average data are summarized in **Figure 8B**. NAbs were detected mainly in the lower dilutions of serum (1/10 and 1/40). For T1, the higher levels of neutralization were observed for G1 and G3. However, no significant differences were observed for serum samples collected at T2 after vaccination. In T3, G1 and G2 showed slightly greater levels of neutralization on average. Before vaccination (T1), 19.5%, 9.8% and 16.3% of individuals had NAb titers ≥1/40 (considered as protection) for G1, G2 and G3, respectively (**Figure 8C**). However, after vaccination, the percentage of individuals showing protective NAb titers (≥1/40) was 17.1% (T2) or 19.5% (T3) for G1, 31.4% (T2) or 11.8% (T3) for G2, and 39.5% (T2) or 6.98% (T3) for G3 (**Figure 8C**). Although most of individuals only displayed protective titers of NAbs (≥1/40) in the dilution 1/40 (**Figure 8B**). The data suggest that seasonal IIV contributed to increase the levels of NAbs in all groups, although this effect was more evident for G2 and G3, which could be associated with immunosenescence in G1. Crystal violet staining, although is a more subjective and less quantifiable method, confirmed the results obtained when Nluc activity was measured in the tissue culture supernatant (data not shown). These results in the MN assays, although similar, did not correlate perfectly with those of the HAI assay, but this was expected given the different nature of these assays (35, 43). HAI assays are only able to detect RBD HA NAbs while the MN assay are able to detect RBD and stalk HA, and NA NAbs (38, 43).

**Figure 8.**
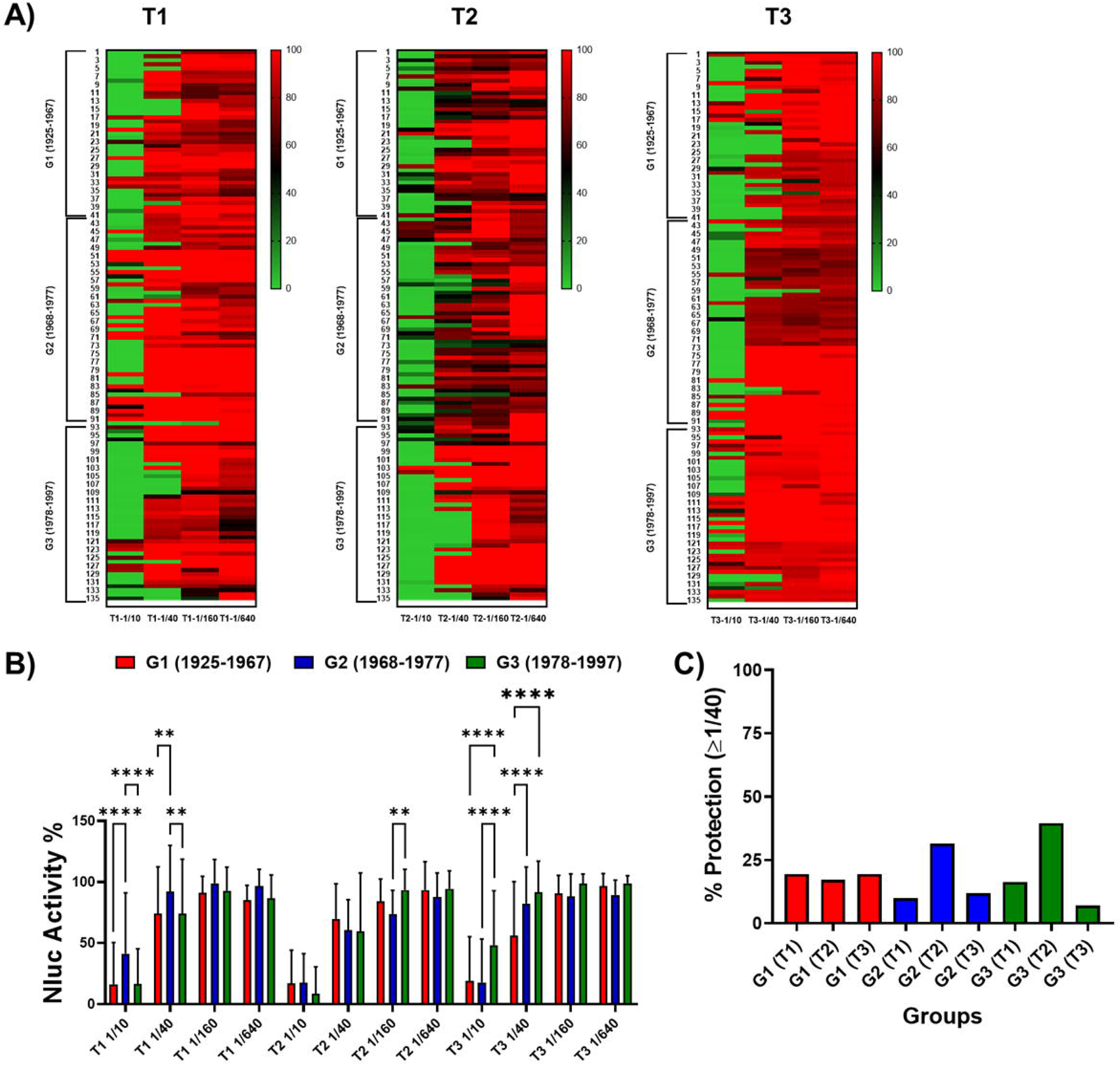
Identification of cross-neutralizing antibodies against H5N1. One-two hundred FFU of PR8(H5N1)-Nluc was preincubated for 1 h with 4-fold serial dilutions (starting dilution 1/10) of the human serum samples corresponding to individuals born 1925-1967 (G1), 1968-1977 (G2), and 1978-1997 (G3) and collected immediately before vaccine administration (T1), and post-vaccination at 28 days after vaccination (T2) or six months after vaccination (T3). Subsequently, MDCK cells (96-well plates, 2 x 10^4^ cells/well, and triplicates) were infected with the antibody-virus mixture. Virus neutralization was determined by quantitating NLuc reporter expression at 48 h p.i.. Mock-infected cells were used as internal controls for basal levels of luciferase (NLuc) expression. Infected cells in the absence of serum were used to determine maximum (100%) reporter expression. Nluc activity or neutralization for the individual serum samples is displayed using a heatmap visualization method (**A**). The average for the different groups is also represented (**B**). *\*p* < 0.05; *\*\*p* > 0.01; *\*\*\*p* < 0.001; *\*\*\*\*p* < 0.0001, using two-way ANOVA. Red indicates more Nluc activity or less neutralization, and green indicates less Nluc activity or more neutralization. **C**) The percentage of individuals showing protection (NAb titers ≥1/40) is also presented.

### 3.5 Cross-reactive antibodies to neutralize IAV H7N9 in vitro

The presence of NAbs against IAV H7N9 was also evaluated for the 135 individuals distributed in G1, G2 and G3, before vaccination (T1) or after vaccination (T2 and T3). For that, Nluc-based MN assays were performed using the same human serum samples and PR8(H7N9)-Nluc (**Figure 9**). The data suggest that, at T1, G1 shows higher levels of NAbs. In contrast, at T2, groups G2 and G3 showed higher levels of NAbs. The levels of NAbs at T3 were similar to those detected in T1 for the 3 age groups (G1, G2 and G3) (**Figure 9A and B**). As in the case of NAbs against H5N1, most of individuals had detected NAbs only in the lower dilutions of serum (1/10 and 1/40). **Figure 9C** shows the percentages of subjects whose NAb titers were ≥1/40 (considered protection) for each group. These percentages indicate that before (T1) or after vaccination, the NAb titers protected against PR8(H7N9)-Nluc were 29.3% (T1), 12.2% (T2) or 24.39% (T3) for G1; 13.7% (T1), 35.3% (T2) or 15.69% (T3) for G2; and 11.6% (T1), 51.2% (T2) or 16.28% (T3) for G3. Taken together, these data suggest that some weak heterotypic responses to H7N9 virus are induced after vaccination (T2) with seasonal IIV and that the responses are better for G2 and G3. Crystal violet staining confirmed the results obtained in the Nluc-microneutralization assay (data not shown). Similar results were observed in the HAI assays, although with some expected discrepancies as indicated above for H5N1.

**Figure 9.**
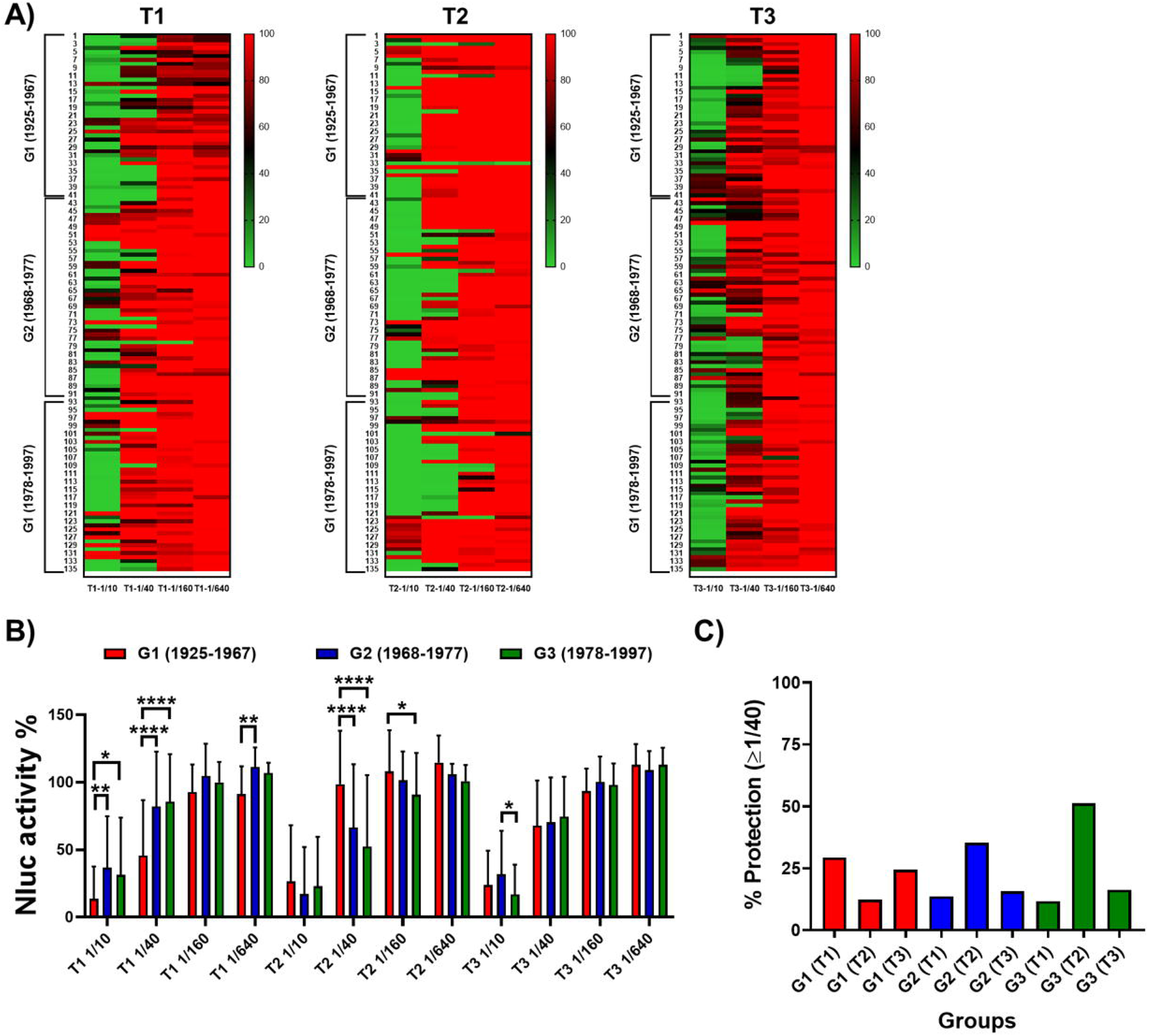
Identification of cross-neutralizing antibodies against H7N9. One-two hundred FFU of PR8(H7N9)-Nluc was preincubated for 1 h with 4-fold serial dilutions (starting dilution 1/10) of the human serum samples corresponding to individuals born 1925-1967 (G1), 1968-1977 (G2), and 1978-1997 (G3) and collected immediately before vaccine administration (T1), and post-vaccination at 28 days after vaccination (T2) or six months after vaccination (T3). Subsequently, MDCK cells (96-well plates, 2 x 10^4^ cells/well, and triplicates) were infected with the antibody-virus mixture. Virus neutralization was determined by quantitating NLuc reporter expression at 48 h p.i.. Mock-infected cells were used as internal controls for basal levels of luciferase (NLuc) expression. Infected cells in the absence of serum were used to determine maximum (100%) reporter expression. Nluc activity or neutralization for the individual sera samples was displayed using a heatmap visualization method (**A**). The average for the different groups is also represented (**B**). *\*p* < 0.05; *\*\*p* > 0.01; *\*\*\*p* < 0.001; *\*\*\*\*p* < 0.0001, using two-way ANOVA. Red indicates more Nluc activity or less neutralization, and green indicates less Nluc activity or more neutralization. **C**) The percentage of individuals showing protection (NAb titers ≥1/40) is also presented.

## 4 Discussion

The threat of AIV, mainly HPAI H5N1, to cause the next influenza pandemic remains a major health concern. In this work, the basal heterosubtypic immunity against H5N1 and H7N9 IAV subtypes was evaluated. In addition, cross-protection induced by human seasonal IIV was determined. The analysis was performed in 3 age groups of adults using approximately 400 serum samples. To perform this study, we engineered four new recombinant viruses expressing the HA and NA of H5N1 or H7N9 strains in the backbone of PR8, including Nluc-expressing versions, to perform Nluc-based MN assays. The identity and replication properties of these viruses were characterized.

It is assumed that a 1:40 titer of antibodies with HAI activity will reduce the risk of being infected with seasonal IAV H1N1 or H3N2 by 50% (38). Although, we are aware that the same may not apply to infections with potentially pandemic or zoonotic IAVs, or to all age groups, for our study we consider this value to be indicative of protection for both HAI and MN assays. Our HAI data, using PR8(H5N1) or PR8(H7N9) inactivated viruses, show limited levels of seroprotection and seroconversion for H5N1 or H7N9 viruses. For H5N1, seroprotection was greater after vaccination (T2) for G2 and G3. However, for H7N9, the percentage of seroprotection was similar in all cases.

Previous results from animal studies or epidemiologic reports indicate that heterosubtypic immunity should be considered as a significant factor in the immune response against novel IAV infections (44). Our MN assay results showed that some basal heterosubtypic immunity against H5N1 and H7N9 IAV subtypes may exist, although at low levels. Moreover, our findings suggest that vaccination against seasonal IAV could boost heterosubtypic immunity against unrelated H5N1 or H7N9 strains. However, despite of the level of NAbs detected, it remains unclear whether heterosubtypic immunity could be correlated with protection efficacy and help to reduce the severity caused by H5N1 or H7N9 IAV infections. In a previously published study by of our group, we demonstrated also a moderate response of heterosubtypic antibodies after seasonal vaccination (45). In this work, we showed that a seasonal IIV is able to induce protective titers against the H5N1 subtype in 20.0-27.9% of the population regardless of their age, but we did not demonstrate an increase in antibodies against H7N9 (45).

Heterosubtypic immunity is a well-described phenomenon for human IAV (44–47). Several clinical studies for H5N1 vaccines have confirmed the presence of serum antibodies against H5N1 in a reduced number of samples collected prior to vaccination (48–52). Recently, van Maurik et al. demonstrated that vaccination of mice with seasonal IIV elicited a humoral heterosubtypic immune responses against H5N1 (44). The authors showed that H1 HA-specific, rather than H3 HA- or B HA-specific IgG antibodies had heterosubtypic reactivity against recombinant H5. This was not surprising since the H1 subtype is more antigenically related to the H5 subtype than to the H3 subtype (53, 54). In another study, groups of ferrets were infected with seasonal IAV H1N1 or H3N2 or immunized with seasonal IIV. Then, serological (HAI or MN) assays revealed that cross-reactivity against the H5N1 virus was not detected in any group (55). Following challenge with A/Vietnam/1203/2004 (H5N1), only infected groups exhibited attenuated viral loads leading to 100% survival. These data suggest that natural infection with human seasonal strains could potentially provide better heterosubtypic protection against HPAI H5N1 virus infection compared to vaccination with IIV (55). In a recent study by Levine et al, it was determined that the population in the United States was immunologically naive to H7N9 HA (56). However, seasonal vaccination induced cross-reactive functional antibodies to H7N9 NA, but minimal cross-reactive antibody-dependent cell-mediated cytotoxicity (ADCC) antibodies. In another study, it was shown that H7 cross-reactive antibodies induced by vaccination with H1N1 or H3N2 strains are not uncommon (57). Three H3 HA-reactive MAbs generated by individuals after IAV vaccination were able to neutralize H7N9 viruses and protect mice against homologous challenge. Interestingly, the H7N9-NAbs bound to the HA stalk domain.

Our results are in concordance with those previous studies, indicating that seasonal IIV could be in some way useful in the case of an AIV pandemic. However, it is important to note that our results also demonstrate that, far from the low-moderate increase in the antibody titers detected only in the younger groups, we also detected that those antibodies were almost completely lost after 6 months. This means that the usefulness of the seasonal vaccines will be likely be limited. However, the results published by Maurik et al. (44) demonstrated that seasonal IIV followed by a H5N1 monovalent vaccine increases both the antibody titers against this specific subtype and also the survival rates of mice challenged with H5N1, compared to a buffer prime followed by H5N1 monovalent vaccination. This is so important since these results could guide us in the vaccination schedule in the case of a H5N1 pandemic. As we observed at the beginning of the COVID-19 pandemic, there was a shortage of vaccine available. Therefore, in the case of an AIV pandemic, the same will likely occur. Currently, there are a total of 175 clinical trials involving AIV pandemic vaccines (https://www.clinicaltrials.gov/search?cond=H5N1%20influenza&term=vaccine&aggFilters=ages:child), but only a few of them (nine) are currently marketed, and most of them are based on old AIV strains. The results of our study and those published by Maurik et al., indicate that probably the best vaccination strategy during the early stages of an AIV pandemic will likely involve immunization first with seasonal IIV followed, when it becomes available, with an AIV-specific vaccine, or with specific newly designed H5N1 vaccines. This would lead to a more robust seroprotection compared with the single administration of the monovalent H5N1 vaccine.

On the other hand, some studies have also demonstrated the importance of the viral NA in heterotypic protection against AIVs. Influenza NA is known to be a major antigenic determinant of IAV and NAbs targeting NA can be induced after IAV infection or seasonal vaccination (47). A study published in 2008 demonstrated that the heterotypic cellular response against H5N1 virus elicited by seasonal vaccines relies mostly on the N1 related to the T-CD4 response (46). Therefore, whether IIV (or viral infection) is able to induce heterosubtypic NA-reactive antibodies against IAV H5N1 or H7N9 should be studied. Therefore, inhibition of NA activity assays could also be important for determining the degree of protection due to anti-NA antibodies. However, the amount of NA contained in IIV is very limited, and IIV induced limited, if any, responses to viral NA (6, 26, 42, 47). In this regard, H5N1 and the H1N1 present in the IIV share similar N1 while this is not the case for H7N9. On the other hand, antibodies targeting the conserved stalk domain of HA could also be involved in the neutralization observed in our assays. However, the mechanisms underlying H5N1 or H7N9 neutralization need to be better characterized. These alternative mechanisms of viral neutralization, could explain the discrepancies observed for some HAI and MN results of our and other studies. However, all these factors were not addressed in this work, but it will be important to evaluate them in future studies. Importantly, T-cell responses may also be key in mediating protection against novel IAV strains, including the H5N1 and H7N9 subtypes (22, 23, 42). Compared with IIV, live-attenuated influenza vaccines (LAIV) have been shown to be more effective at inducing cellular responses and protecting against heterologous or heterosubtipic strains than IIV. Therefore, it could be highly relevant to evaluate the role of LAIV in inducing NAbs against circulating IAV strains with pandemic potential, such as H5N1 and H7N9 viruses.

One weakness of our study is that we used old correlates of protection to evaluate seroprotection (and neutralization), based on the 1:40 seroprotection titer, and that these correlates are not specifically intended for AIV. Additionally, this correlation is specific for HAI, but we also used it for neutralization assays for the comparisons made in this work.

In summary, the immune protection against H5N1 and H7N9 AIV in the population in our study is practically nil since a low percentage of individuals showed protective heterotypic antibodies before vaccination. However, seasonal influenza vaccines elicited seroprotective responses against the H5N1 subtype in nearly 15% of the younger individuals included in the study, indicating that this kind of vaccines could be somehow useful in the case of an H5N1 pandemic. This is important since in the first stages of an AIV pandemic, there will likely be a shortage of specific H5N1 vaccines; therefore, seasonal influenza vaccines could be the first line of defense against the virus and need to be completed after specific H5N1 vaccines are available.

## 6 Author Contributions

Conceptualization, I.S-M., A.N, and J.M.E, Methodology, I.S-M., A.N, A.M., C.S.C-C, J.S-M and C.R-C; Formal Analysis, I.S-M., A.N, C.S.C-C, J.S-M and C.R-C; Funding Acquisition, I.S-M., A.N. and J.M.E; Investigation, I.S-M., A.N, C.S.C-C, J.S-M and C.R-C; Writing – Original Draft: I.S-M., and A.N; Writing-Review & Editing: I.S-M., A.N, J.S-M, C.R-C, C.S.C-C, M.H, S.R-R, M. D-G, A.M, L.M-S, and J.M.E;, and S.M.T. All authors have approved the manuscript for publication.

## 8 Acknowledgments

We thank the Biodefense and Emerging Infectious Research Resources Repository (BEI Resources) for providing the following reagents: NR-2705, NR-665, NR-48597, NR-9598, NR-49276, NR- 59424 and NR-44365.

